# Delayed lubricin injection improves cartilage repair tissue quality in an in vivo rabbit osteochondral defect model

**DOI:** 10.1101/2025.02.06.636825

**Authors:** Donghwan Yoon, Karan Vishwanath, John Dankert, James J. Butler, Mohammad T. Azam, Arianna L. Gianakos, Marshall J. Colville, Serafina G. Lopez, Matthew J. Paszek, Heidi L. Reesink, John G. Kennedy, Lawrence J. Bonassar, Rebecca M. Irwin

## Abstract

Osteochondral lesions (OCL) are common among young patients and often require surgical interventions since cartilage has a poor capacity for self-repair. Bone marrow stimulation (BMS) has been used clinically for decades to treat OCLs, however a persisting challenge with BMS and other cartilage repair strategies is the inferior quality of the resulting fibrocartilaginous repair tissue. Lubrication-based therapies have the potential to improve the quality of cartilage repair tissue as joint lubrication is linked to local cartilage tissue strains and subsequent cellular responses including death and apoptosis. Recently, a full length recombinant human lubricin (rhLubricin) was developed and has been shown to lower friction in cartilage. This study investigated the effect of a single delayed injection of rhLubricin on cartilage repair in an *in vivo* rabbit OCL model using gross macroscopic evaluation, surface profilometry, histology, and tribology. Moderate improvement in macroscopic scores for cartilage repair were observed. Notably, quantitative analysis of Safranin-O histology showed that rhLubricin treated joints had significantly higher glycosaminoglycan content compared to saline treated joints, and there were no differences in repair integration between groups. Furthermore, rhLubricin treated joints had significantly lower friction coefficients tested across three sliding speeds compared to saline treated joints (rhLubricin: 0.15 ± 0.03 at 0.1 mm/s to 0.12 ± 0.03 at 10 mm/s, Saline: 0.22 ± 0.06 at 0.1 mm/s to 0.19 ± 0.05 at 10 mm/s). Overall, a single delayed injection of rhLubricin improved the quality and lubricating ability of the repair cartilage tissue without inhibiting repair tissue integration.

## 1 Introduction

Osteochondral lesions (OCLs) are typically the result of a traumatic injury to cartilage and/or the underlying bone in joints. They are common in the knee and ankle joint and may be associated with chondral degeneration and the development of osteoarthritis.^1^ As cartilage has a poor capacity for self-repair, surgical intervention is often needed.^2,3^ Bone marrow stimulation (BMS) is the current gold standard repair method that involves creating a bleeding bed by violating the subchondral plate thereby releasing mesenchymal stem cells and promoting hyaline-like tissue repair.^4^ However, BMS produces fibrocartilage that is mechanically inferior compared to native cartilage tissue and therefore leaves the joint susceptible to mechanically induced damage.^5–8^ OCLs are prevalent in young adult patients, highlighting the clinical need for surgical interventions that restore the mechanical function of cartilage tissue while preserving joint health. ^9,10^

Microscale assessment of the mechanical strains and stresses as well as tribological properties of healthy and diseased tissue in several species have been well documented in cartilage explants.^11–16^ Specifically, microscale tissue strains strongly correlate with cellular outcomes where increased tissue strains lead to increased cell death, apoptosis, and mitochondrial dysfunction in cartilage tissue.^13,17–20^ Numerous studies have shown that alterations in lubrication change cartilage tissue strains, and subsequently will affect cell responses.^21–23^ Cartilage lubrication is mainly facilitated by two macromolecules in the synovial fluid (SF): hyaluronic acid and lubricin.^24–26^ In order to create a lower stress environment in articulating joints, viscosupplementation, or supplementation with a highly viscous material like hyaluronic acid, has been the cornerstone of OA treatment for decades with variable success.^27,28^ Recently, a bioengineering approach using codon optimization was used to produce a recombinant lubricin with the native human N- and C-terminal domains and a lubricin-inspired mucin domain containing 59 KEPAPTTP-rich repeats (rhLubricin).^29,30^ Lubricin’s role in cartilage lubrication has been well documented with several studies reporting increased friction with lubricin degradation and decreased friction with lubricin supplementation across various tribometer systems.^23,30,31^ Lubricin has also been purported to have an anti-inflammatory role and could possibly lower the interfacial strains in the joint.^18,32^ However, little is known about its ability to alter the quality of repaired cartilage.

Previously, it has been shown that lubricin incubation performed immediately after OCL defects inhibited integration between repair and native cartilage due to its anti-adhesive properties in vitro.^33^ However, lubricin is critical for proper joint lubrication, maintaining low friction in the joint and preserving cell viability.^18^ Therefore, we hypothesized that by delaying the administration of lubricin into the joint following OCL repair such that repair tissue has already filled in the defect site, lubricin will improve the quality of the repair tissue without impeding the integration of the repair and adjacent native tissue. To investigate this hypothesis, this study evaluated whether a single delayed injection of rhLubricin can improve cartilage repair in a New Zealand rabbit OCL model. We hypothesize that rhLubricin injections will: (A) improve the quality of the repair tissue, (B) improve the mechanics of the repair tissue, and (C) not impede the integration of the repair and native cartilage tissue.

## 2 Materials and Methods

### 2.1 In vivo study design

The NYU Institutional Animal Care and Use Committee approved all surgical procedures and experimental designs (IACUC 2017-0025). Bilateral 3 mm diameter osteochondral (OCL) defects along with bone marrow stimulation (BMS) were performed on the medial femoral condyle of 16 female New Zealand rabbits. A recombinant full-length human lubricin was produced and purified as previous described.^30,34^ A saline vehicle control was formulated as 20 mM Na_2_PO_4_, 100 mM NaCl, pH 7.2 to match the rhLubricin solution (Sigma-Aldrich, St. Louis, MO). Both rhLubricin and the vehicle were terminally sterilized by 0.22 μm filtration (Pall Corporation, New York, NY), aliquoted into septum-top serum vials, frozen in liquid nitrogen and stored at −80 °C until use. The timepoint of rhLubricin and saline injections were performed 4 weeks after surgery, thereby providing sufficient time for repair tissue infill to have been achieved.^6^ Injection volumes were 100 μl for saline and rhLubricin. Joints were randomized for treatment and injections were performed in an unblinded manner. The concentration of rhLubricin was 1 mg/ml, which was shown to be within the same order of magnitude of friction data for other forms of recombinant lubricin.^23,30^ Control limbs received no defect and no injection. Confounders such as the order of treatments and measurements were not controlled for in this study. Animals were euthanized at 12 weeks post-operatively, limbs were stored on ice, and the femoral condyles were collected within 12 hours (Figure 1). A total of 18 rabbits (36 joints) were randomly separated into three groups: rhLubricin n = 12 joints; saline n = 16 joints; control n = 8 joints. One saline joint was excluded from the analysis due to an exposed joint capsule. Joints were labeled in 50 ml falcon tubes (VWR International, Radnor, PA) with sample numbers (blinded by DY and KV) macroscopically assessed, analyzed using profilometry and subsequently divided for friction testing or histological analyses.

**Figure 1.**
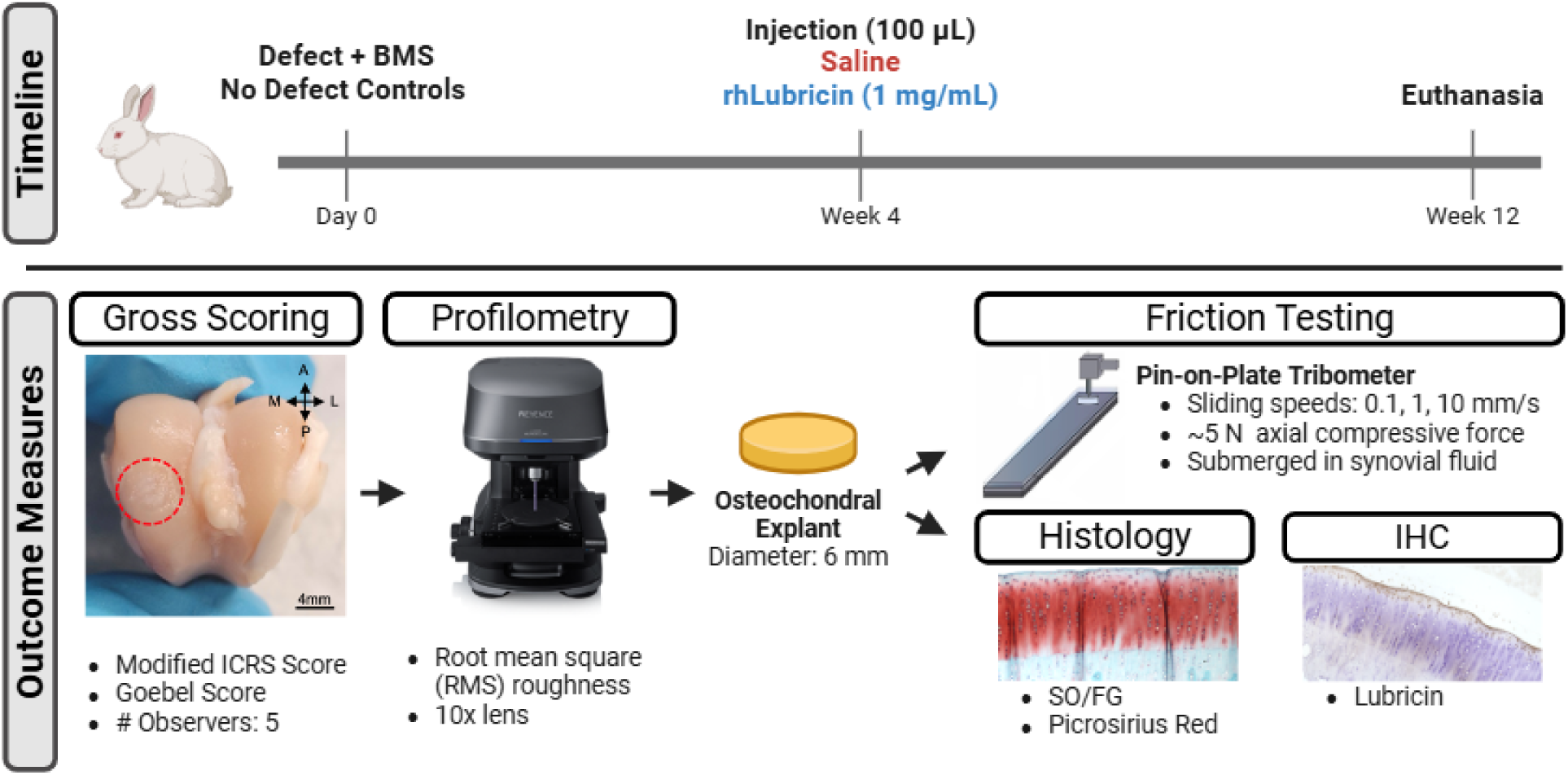
Flowchart of study design and methods used to investigate the outcome measures of gross scoring, surface roughness, coefficient of friction, histology, and lubricin immunohistochemistry of the study.

### 2.2 Macroscopic evaluation of repair tissue

To assess the quality of the repair, macroscopic evaluation was performed. Modified International cartilage regeneration repair assessment (ICRS) scores^35,36^ and Goebel scores^37^ were collected from 4 - 5 blinded observers for the saline and rhLubricin groups (n = 6) on the OCL repair site of the joint. Observers were given a visual training set to become accustomed to the scoring criteria (Table S1, 2). The two methods were both evaluated and median scores from the 4 - 5 observers were compared between saline treated and rhLubricin treated groups.

### 2.3 Surface profilometry

To assess the roughness and surface characteristics of the repair tissue, the root mean square surface roughness (*R_q_*) was measured in the central region of the defect site on the medial femoral condyle (rhLubricin n = 10, Saline n = 15, Ctrl n = 6) using a Keyence VK-X260 confocal laser-scanning profilometer.^38,39^ Roughness measurements were taken with a 10x lens that has a working distance of 16.5 mm and capture size of 1412 μm x 1060 μm. The laser-scanning profilometer scans a convergent laser beam over the surface of the tissue. No physical interaction was present, therefore allowing the surface of the tissue to remain intact and hydrated to minimize the effect of surface degradation or dehydration for further analyses. The data was smoothed by fitting the contour of the cartilage surface to a 5-parameter waveform. Root mean square (RMS) roughness was measured in the center of the defect with a vertical and horizontal measurement. The average RMS roughness was calculated and reported as the roughness of the central region of the defect site.

### 2.4 Sample preparation

Following macroscopic scoring and profilometry, full thickness 6 mm diameter cylindrical osteochondral explants were obtained from defect site of the medial femoral condyles of the rabbits. The cylindrical explants of the control, saline, and rhLubricin treated groups were divided into two groups: (1) friction testing or (2) histological staining. Samples designated for histological staining were bisected into hemicylinders in the posteroanterior direction. The bisected hemicylinders were frozen immediately after collection and were stored without any buffer to prevent crystallization forming on the cartilage.

### 2.5 Histology, GAG quantification, and histopathology scoring

Histological staining of the samples (n = 3 per group) was performed by the Clinical Pathology Core (Cornell College of Veterinary Medicine, Ithaca, NY). The bisected hemicylinder explants were fixed in 10 v/v% neutral buffer formalin for 24 - 48 hours, rinsed with 70% ethanol, decalcified in a solution made from equal parts of 20% Sodium Citrate and 44% Formic Acid, sectioned in 4 μm thickness slices, and stained with Safranin-O and Picrosirius red to observe relative proteoglycan content and collagen content, respectively. The sections were imaged using a Keyence BZ-X810 microscope at 10x magnification. Proteoglycan to defect area ratio was quantified using ImageJ (NIH, Bethesda, MD) by measuring the area of the 3 mm wide defect and manually quantifying the area of proteoglycan content within the 3 mm wide defect area.^40^ The manual area coverage quantification was repeated five times and averaged to reduce variation within measurements. To assess the histopathological grading of the repair tissue, consensus scoring was performed by three observers (EY, KV, RMI) using a modified version of a previously established semi-quantitative scoring system was used (Table S3).^41^ Observers were blinded to treatment for scoring.

### 2.6 Immunohistochemistry

Lubricin immunohistochemistry was performed on osteochondral sections from the rabbit condyles to further investigate lubricin content and distribution in these samples as previously described.^42^ Samples were fixed, decalcified, paraffin embedded, and sectioned as described above (*Histology, GAG quantification, and histopathology scoring* section). Immunohistochemistry was conducted using a modified Vectastain ABC kit protocol (Vector Laboratories, Burlingame, CA) and an anti-lubricin monocloncal antibody 9G3 at a 1:250 dilution (MABT401, Sigma Aldrich, St. Louis, MO).^42^ Controls were run in parallel with samples and control and experimental samples were stained in the same batch processes and exposed to the same duration and concentration of reagents. All slides were counterstained with hematoxylin. Images were obtained with a Keyence BZ-X810 microscope at 10x magnification. Images were stitched with ImageJ (NIH, Bethesda, MD).^40^

### 2.7 Tribological evaluation of the osteochondral cores

Tribological characterization of the osteochondral cores (rhLubricin n = 4, Saline n = 6, Ctrl n = 6) was performed using a previously described, custom cartilage-on-glass tribometer.^22,23^ Frozen osteochondral cores from all treatment groups were thawed in PBS with protease inhibitors (50ml buffer per tablet, Cat. No. A32965, Thermo Fisher Scientific, Waltham, MA) prior to the friction tests. The samples were glued to brass posts (6 mm diameter), mated against a polished glass surface and tested in healthy bovine synovial fluid (BSF, Lampire Biologics, Pipersville, PA). The cores were then compressed to a normal force of 5.5 ± 1.5 N and allowed to stress relax for 1 hour. After equilibration, the glass counter-face was reciprocated at sliding speeds of 0.1, 1, and 10 mm/s using a DC motor. These compression levels and sliding speeds were chosen based on the strong correlation of the reported friction data to clinical outcomes and biochemical markers of inflammation.^44^ The coefficient of friction (µ) was recorded as the ratio of shear to normal force measured by a biaxial load cell. All friction coefficients were calculated at the end of sliding when friction reached an equilibrium value and then averaged in the forward and reverse sliding directions to give a mean value for the coefficient of friction at each speed.^43^

### 2.9 Statistical Analyses

#### 2.9.1 Macroscopic and histological scoring for cartilage repair

Differences in the macroscopic scoring for cartilage repair between the rhLubricin and saline treated samples and statistical significance for histological scoring and the GAG/defect area ratio was determined using Tukey’s Honestly Significant Difference (HSD) test in R Studio (Posit Software, Boston, MA). Significance was evaluated at *p* < 0.05.

#### 2.9.2 Surface roughness data

Differences in the RMS roughness between the lubricin and saline treated samples were evaluated using Welch’s two-sample t-test using the t-test function in R Studio (Posit Software, Boston, MA). Significance was evaluated at *p* < 0.05.

#### 2.9.3 Tribology data

A linear mixed effects regression model was used to fit coefficient of friction as a function of the treatment at each speed. Random effects in the model included the limb nested within each rabbit. *Post-hoc* pairwise comparisons (adjusted for multiple groups) were conducted to estimate the difference in the estimated marginal means of the coefficient of friction between the rhLubricin, saline, and control groups using the emmeans package in R Studio. Statistical significance was evaluated at *p* < 0.05. A repeated measures Pearson correlation analysis was conducted in R Studio using the rmcorr and lme4 packages to assess the difference between the coefficient of friction of the cores for the three groups, with the articulation speeds used as the repeated measure in the analysis.

## 3 Results

### 3.1 Gross repair tissue characterization

Gross images of the OCL defects demonstrated repair tissue infill following rhLubricin or saline injections (Fig. 2A). Visual examination showed the repair tissue was more fibrillated and rougher compared to the surrounding native tissue for both treatments. For further quantitative analysis, gross macroscopic scoring of the repair tissue using a modified ICRS score, and a Goebel score was performed (Fig. 2 B, C). The scores for healthy control condyles were 16 and 20 for the modified ICRS score and Goebel score, respectively. The median values for saline and rhLubricin for the modified ICRS score were 7.8 and 10.3, respectively. The median values for saline and rhLubricin for the Goebel score were 10.5 and 13.5, respectively. Both scoring methods showed a trend for improved repair with rhLubricin compared to saline but were not statistically significant (ICRS: p = 0.08, Goebel: p = 0.11). Notably, there were no differences between saline and rhLubricin treated joints in the subscores that corresponded to repair tissue integration (Fig. 2C, E, *p* > 0.90). The remaining subscores for each scoring system are presented in Supplemental Tables 4 and 5.

**Figure 2.**
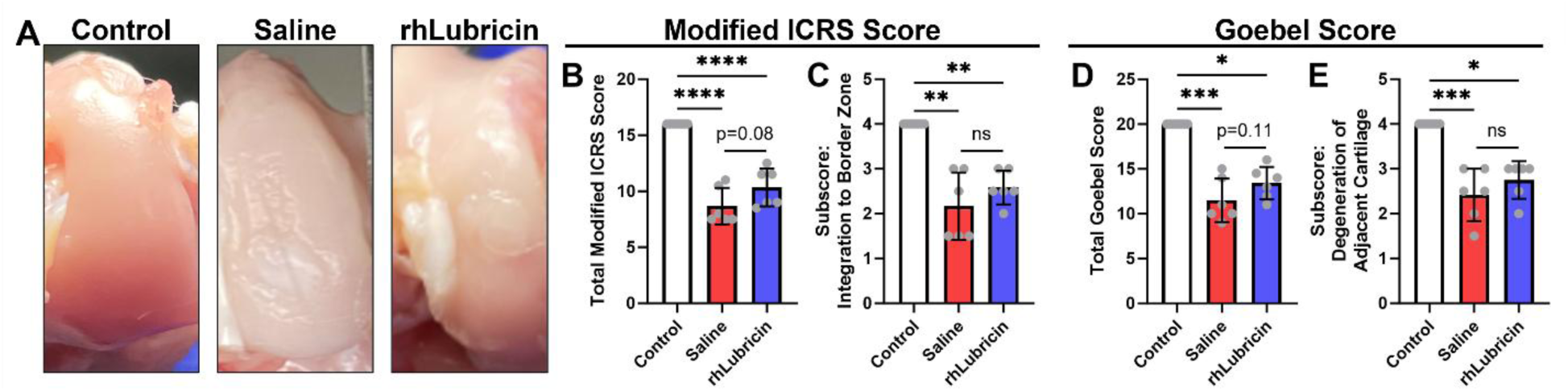
Gross image of the medial condyle with repair site of the defect area highlighted, with gross scoring of the condyle using a modified ICRS and Goebel score. Subscores pertaining to repair tissue integration are shown for both the ICRS (C) and Goebel (E) scores. Asterisks indicate statistical significance (\**p* < 0.05, \*\**p* < 0.01, \*\*\**p* < 0.001, \*\*\*\**p* < 0.0001).

### 3.2 Histological assessment of tissue repair quality and integration

Histological staining revealed spatial distributions of proteoglycans, lubricin, and fibrillar collagen in the control and repair tissue samples. Both defect groups showed decreased proteoglycan content in the repair tissue compared to adjacent healthy cartilage and to the uninjured control (Fig. 3 A - C). Within the defect groups, joints treated with rhLubricin had increased proteoglycan staining in the deeper region of repair tissue compared to saline treated joints (Fig. 3 B, C). In the uninjured control and the cartilage tissue adjacent to the defect for both saline and rhLubricin treated joints, lubricin staining was concentrated at the surface of the cartilage tissue (Fig. 3 D (i), E (ii), F (iv)). rhLubricin treated joints had enriched lubricin staining at the tissue surface compared to saline treated joints in the cartilage tissue adjacent to the defect (Fig. 3 E (ii), F (iv)). For the repair cartilage tissue, saline treated tissue had lubricin staining throughout with no apparent differences through the repair tissue depth (Fig. 3 (iii)). In contrast, the rhLubricin repair tissue had enriched lubricin staining at the surface of the tissue that diminished in the deeper regions (Fig. 3 (v)). Notably, there was no enrichment of lubricin staining at the interface of repair for either saline or rhLubricin treated groups. Additionally, there were no obvious differences in collagen organization observed under brightfield and polarized light between saline and rhLubricin groups (Fig. 3 J - L).

**Figure 3.**
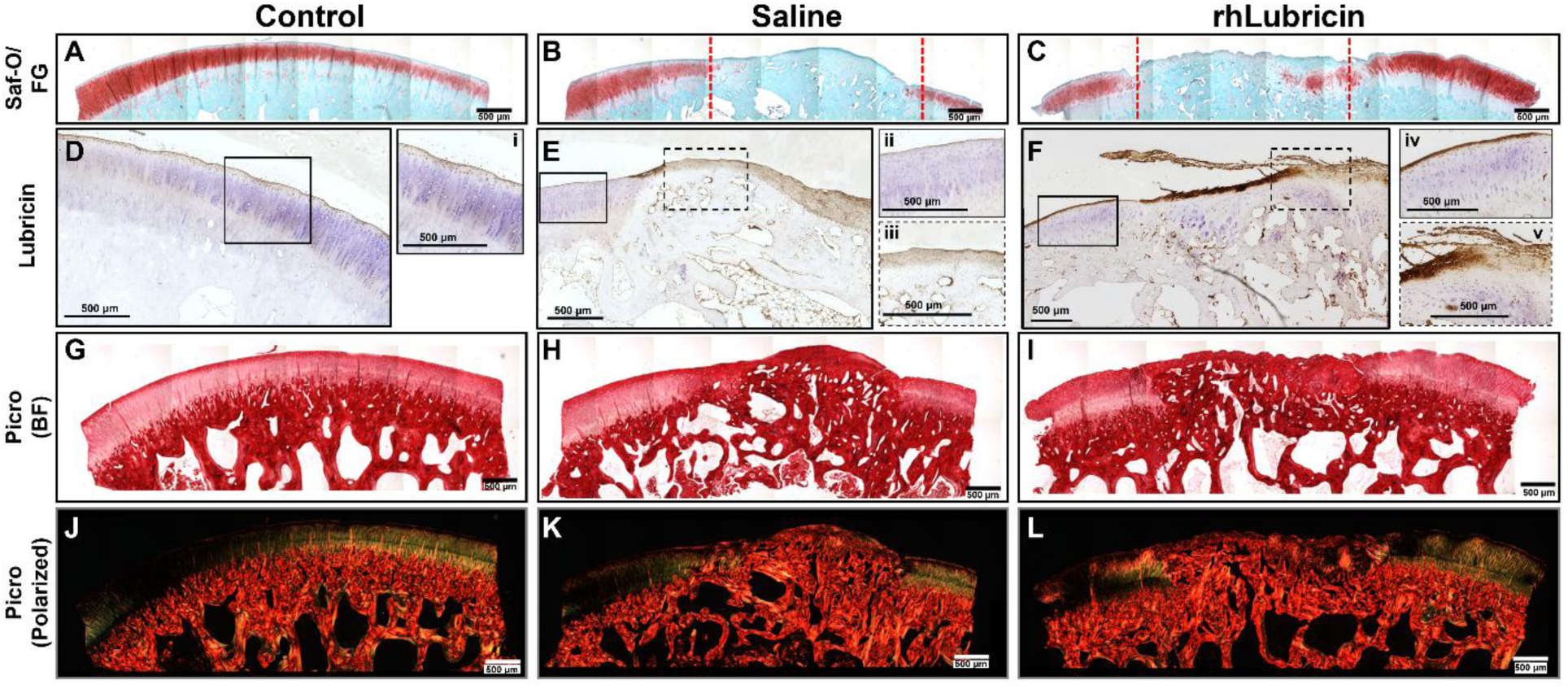
Histological stains of Safranin-O, lubricin immunohistochemistry, and Picrosirius red observed under brightfield (BF) and polarized light. Dashed red lines in panels B and C indicate the location of the defect. Scale bar = 500 μm for all images.

An enrichment of GAG content in the rhLubricin treated group relative to the saline treated group was observed by histological staining with Safranin-O. This observation was quantified using a GAG to defect area (3 mm) ratio with mean values of 63 %, 32 %, and 12 % for control, rhLubricin, and saline respectively (Fig. 4A). The rhLubricin treated group did not restore the GAG content to native levels (*p* < 0.05). However, the GAG/Defect area ratio demonstrated that rhLubricin treated group had a significant improvement in GAG content compared to the saline treated group (*p* < 0.05). Histological assessment performed using a previously established scoring system (Table S1)^41^ showed differences between healthy and defect tissue (Fig. 4B - J). Combined histopathological scoring indicated there was a clear difference between healthy controls and the rhLubricin and saline treated defect groups (Fig. 4B). Individual sub-scores of cell distribution, Safranin-O stain, and tidemark reformation depicted improvements in the quality of repair for the rhLubricin treated group compared to the saline treated group (Fig. 4C, I, J). Notably, there was no difference in tissue integration scoring between saline and rhLubricin treated groups (Fig. 4E). Safranin-O stained sections used for histological scoring can be found in Supplemental Figure 1.

**Figure 4.**
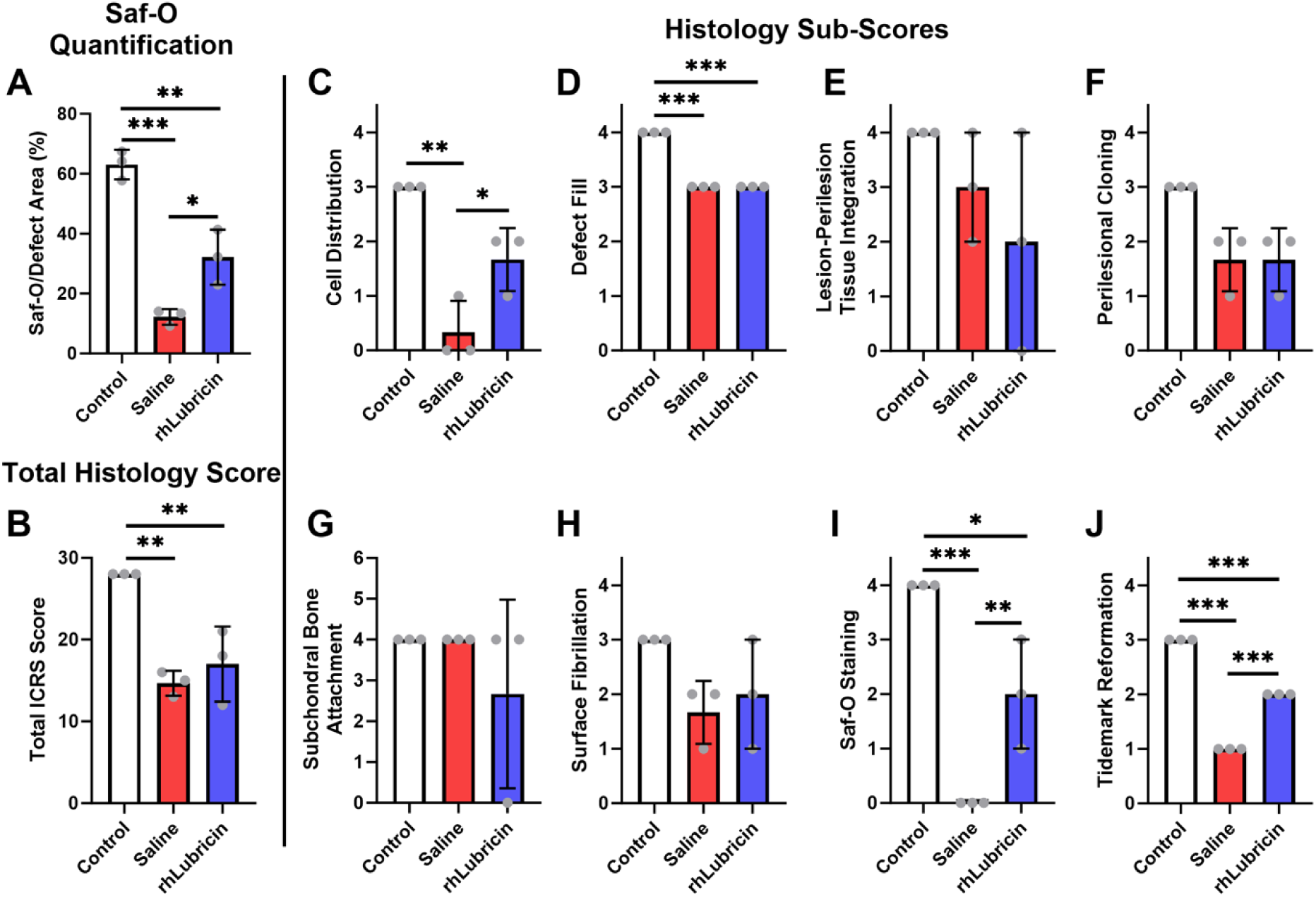
Quantification of the Safranin-O stains to defect area and histological grading of total and sub-scores. Asterisks indicate statistical significance (* *p* < 0.05, ** *p* < 0.01, *** *p* < 0.001).

### 3.3 Surface roughness and tribological evaluation

Surface profilometry revealed differences between the roughness of the center region of the control and repair tissue for both rhLubricin and saline treated groups. The mean value for the RMS roughness in the center of the defect site for the healthy control condyles was 4.0 ± 1.3 µm. The rhLubricin treated group had lower surface roughness measurements compared to the saline treated group, but this was not statistically significant (*p* = 0.16; rhLubricin: 8.1 ± 2.7 µm; saline: 11.5 ± 7.2 µm). Surface roughness measurements for rhLubricin and saline treated samples were both significantly higher than the healthy control (*p* < 0.05) (Fig.5 A).

**Figure 5.**
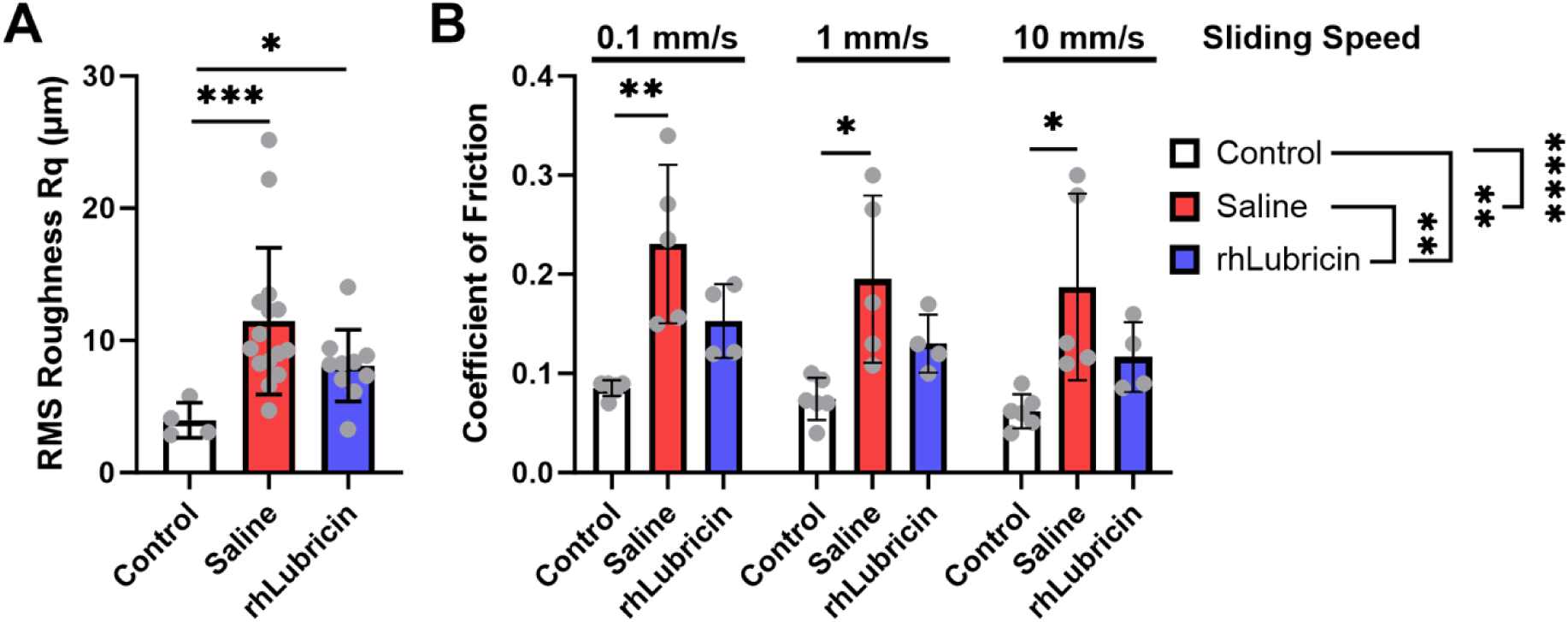
Surface roughness measurements and coefficient of friction of the center of the defect. Treatment comparisons within sliding speeds are from a post-hoc analysis from a linear mixed effects model. Treatment level statistical significance reported adjacent to the treatment labels in panel B are from a repeated measures linear mixed effect model where sliding speed was a repeated measure. Asterisks indicate statistical significance (* *p* < 0.05, ** *p* < 0.01, *** *p* < 0.001, **** *p* < 0.0001).

Tribological evaluation revealed distinct differences in friction coefficient of osteochondral cores between the control (no defect), rhLubricin and saline treated groups when lubricated in healthy bovine SF. The coefficient of friction of the control samples was considerably lower than the rhLubricin or saline treated groups across two orders of magnitude of sliding speed (Fig. 5, *p* < 0.05), with friction ranging from µ_avg_ = 0.08 ± 0.01 at 0.1 mm/s to 0.065 ± 0.02 at 10 mm/s. The rhLubricin treated samples had moderate friction coefficients ranging from 0.15 ± 0.03 at 0.1 mm/s to 0.12 ± 0.03 at 10 mm/s, and the saline treated groups had the highest friction coefficients, ranging from 0.22 ± 0.06 at 0.1 mm/s to 0.19 ± 0.05 at 10 mm/s. The control samples had friction coefficients that were significantly lower than the saline treated samples (Fig.5, *p* < 0.05 from 0.1 - 10 mm/s), but were no statistical differences between the control and rhLubricin treated samples (*p* = 0.27, 0.38, 0.49 from 0.1 – 10 mm/s). Additionally, there were no statistically significant differences in the friction coefficient between the rhLubricin and saline treated groups at any given sliding speed (Fig. 5B, *p* > 0.05). However, when comparing the friction coefficient of the overall treatments using sliding speed as repeated measures, rhLubricin treated joints were significantly lower than saline treated controls (Fig.5B, *p* < 0.01).

## 4 Discussion

The objective of this study was to investigate whether a single delayed injection of lubricin after surgery would promote cartilage regeneration in an *in vivo* pre-clinical rabbit model. Our results show that the delayed injection of rhLubricin after allowing initial repair tissue infill improved the overall quality of the repair cartilage tissue. Increased GAG deposition, improved macroscopic histopathological characteristics, and friction coefficients not different from healthy controls were observed in rhLubricin treated repair tissue. Sub-scores for histopathology showed improvements in cell distribution, proteoglycan staining, and tidemark reformation and there was no difference in repair cartilage tissue integration. These results suggest that delayed lubricin injections after OCL repair surgery could provide a therapeutic benefit for overall cartilage repair tissue quality without compromising repair tissue integration.

Lubricin is a mucinous glycoprotein that acts as a boundary lubricant in articulating joints and is critical for joint health.^18^ In mice with a homozygous lubricin knockout (Prg4^-/-^), the whole joint coefficient of friction is increased along with elevated levels of chondrocyte apoptosis, highlighting the importance of lubricin in cartilage lubrication.^18^ A concern for the application of lubricin for repair of cartilage focal defects is the integration of the repair tissue as lubricin coating the surface of both cartilage and meniscus tissues has been shown to inhibit integrative repair.^33,45,46^ Specifically for cartilage, it has been shown that exogenous lubricin supplementation in culture inhibits cartilage repair in vitro.^33^ To prevent the coating of the repair tissue interface with exogenous lubricin, rhLubricin was delayed by 4 weeks after creating an osteochondral defect with BMS. Previous studies have shown with this osteochondral defect model that 4 weeks is sufficient for repair tissue infill into the defect site.^6,47^ Gross evaluation of the repair tissue using the established modified ICRS score and Goebel scoring systems showed that the rhLubricin treated group did not impede the repair tissue integration. Histologically, there was no difference in integration from ICRS scoring and there was no enrichment of lubricin at the repair interface. Thus, lubricin supplementation did not impede or inhibit the integration of repair tissue for OCL defects in this study.

While BMS is considered to be the gold standard for cartilage repair, one of the prevailing challenges is the infill of fibrocartilage repair tissue that lacks the structure, composition, and mechanical properties of native hyaline cartilage.^4^ In this study, augmenting BMS with a delayed lubricin injection resulted in superior repair tissue quality. Increased GAG content was qualitatively observed and quantified in the cartilage repair tissue of rhLubricin treated groups compared to the saline treated group. This coincides with previous studies that saw increased GAG content with viscosupplementation.^48^ As GAGs are critical for the compressive properties of cartilage tissue, this elevated GAG content could likely influence the compressive properties of cartilage as shown in previous studies thus leaving the repair cartilage tissue less prone to mechanically induced damage.^49^

Further histological analysis with a modified version of a previously established semi-quantitative scoring system revealed interesting features between healthy controls, saline-treated defects, and rhLubricin-treated defects. Although combined histological scores within the defect group which were injected with either saline or rhLubricin did not show any significant differences, meaningful differences were noted within individual features that were analyzed.

Apart from the improvement of restoration of GAG content, improvement in cell distribution as well as tidemark reformation were observed within the individual subscores for the rhLubricin group compared to saline treated. Proper cell distribution is critical in to produce the correct extracellular matrix therefore decreasing the probability of compromised mechanical integrity.^50,51^ Improved tidemark reformation in the rhLubricin treated groups also suggests improvement in the long-term stability of the boundary between calcified and non-calcified cartilage which maintains the structure and function of the repair tissue. Incomplete or delayed tidemark reformation also suggests inferior quality of repair which may be prone to degeneration.^52^

The difference in surface roughness measured from profilometry revealed decreased surface roughness for the rhLubricin treated group compared to the saline treated group and was trending towards statistical significance (*p* = 0.16). Control group samples exhibited surface roughness values similar to those previously noted in literature.^53^ Although not statistically different, rhLubricin treated samples were observed to have a mean value of 4 µm of difference in surface roughness compared to saline treated tissue, which corresponds to the total roughness of a healthy condyle. Notably, the coefficient of friction was significantly different between saline and rhLubricin treated groups where the rhLubricin treated joints had significantly lower friction. In OA models, it has been shown that the lubricating properties have a strong correlation to clinical data where lower coefficient of frictions correlated with lower pain scores.^44^ The rhLubricin in the present study lowered friction in repair cartilage tissue, consistent with its ability to lower friction in healthy cartilage explants, and therefore may have implications for affecting patient pain that could be evaluated in future studies.^34^

The influence of lubricin on cartilage health has been studied across multiple animal models. The majority of these studies have been performed in OA models, and several reports have linked lubricin depletion in the knee joint with OA development.^18,33,54–57^ In addition, a recent review found that lubricin supplementation across multiple OA models has shown beneficial effects based on histological, morphological, functional, and gene expression outcome measures.^54^ Notably, studies injecting lubricin supplementation for cartilage defect models are scarce and highlight the importance of this study. Furthermore, the dose of lubricin injection and the content of lubricin varies between models complicating direct comparison of results. As such, this study selected the rhLubricin dose that correlates to coefficients of friction within the same order of magnitude (μ ≅ 0.1) as previously reported for other variants of lubricin.^23,30,58^

There exist several limitations in this study that must be addressed while interpreting these results. First, this study investigated a single injection at a single time point. Future work could address multiple injections at different time points to answer whether increased rhLubricin injections or earlier/later injection time points improves or dampens the outcomes identified from this study. Second, the sample size for this study was determined using a power analysis based on the primary outcome variable, the histological score. This sample size was sufficient to detect statistical differences in the histological and functional lubricating properties of the three groups (control, rhLubricin treated, and saline treated). However, it is likely underpowered to detect smaller effect sizes in secondary outcomes, including gross scoring, surface roughness, and friction coefficients, particularly when comparing rhLubricin to saline or control groups. With an increased sample size, further differences between control tissue, saline treated, and rhLubricin treated repair tissues would likely be observed for gross scoring (ICRS and Goebel) and friction measurements. Future studies should include a larger sample size to properly characterize these differences. Furthermore, this study evaluated a single dose of rhLubricin (100 μl of 1 mg/ml) based off in vitro friction measurements, but other dose concentrations could be explored.^34,44^ While important, these complexities were outside of the scope of this study.

Overall, these data identified improvements in overall quality of repair tissue as well as functional improvements in the lubricating ability of the rhLubricin treated repair cartilage compared to saline treated defects. Additionally, rhLubricin treated joints had enriched lubricin staining at the tissue surface compared to saline treated joints in the cartilage tissue adjacent to the defect. Importantly, there was no difference in repair tissue integration between the saline treated and rhLubricin treated groups. Together, these results identify that a single delayed injection of rhLubricin in an in vivo rabbit model does not inhibit cartilage repair and improves the quality of the repair tissue.

## Acknowledgments

This work was partially funded by the Arthroscopic Association of North America (JGK, RMI), the Cornell Ignite Fellow for New Venture (MJC), the Cornell Technology Acceleration and Maturation Fund (HLR, MJP), the Cornell Technology Acceleration and Maturation Fund (MJC), the NSF LEAP-HI CMMI 2245367 (LJB, HLR, MJP), and the National Institute of Arthritis and Musculoskeletal and Skin Diseases of the National Institutes of Health under Award Number T32AR078751 (SGL). This work made use of the Cornell Center for Materials

Research shared instrumentation facility. The authors would like to acknowledge Caroline Thompson, Ruben Trujillo, Salman Matan, and Megh Prajapti for their assistance with gross scoring of the condyles.

## Author contributions

Research design: JGK, LJB, RMI; acquisition and analysis of the data: DY, KV, JD, JJB, MTA, ALG, MJC, SGL, MJP, HLR, RMI; critical interpretation of results: DY, KV, LJB, RMI; drafted the manuscript: DY, KV, SGL, RMI; read and edited the manuscript: All authors

