## Supplemental for "Delayed lubricin injection improves cartilage repair tissue quality in an in vivo rabbit osteochondral defect model"

**Supplementary Material**

Table S1. Scoring criteria for the modified ICRS score

| | Modified ICRS | Points | |
|---|---|---|---|
| 1 | Degree of defect repair | In level with surrounding tissue | 4 |
| | | 75% repair of defect depth | 3 |
| | | 50% repair of defect depth | 2 |
| | | 25% repair of defect depth | 1 |
| | | 0% repair of defect depth | 0 |
| 2 | Integration to border zone | Complete integration with surrounding cartilage | 4 |
| | | Demarcating border < 1mm | 3 |
| | | 3/4 of graft integrated, 1/4 with a notable border > 1mm width | 2 |
| | | 1/2 of graft integrated, 1/2 with a notable border > 1mm width | 1 |
| | | From no contact to 1/4 of graft integrated with surrounding cartilage | 0 |
| 3 | Macroscopic appearance | Intact smooth surface | 4 |
| | | Fibrillated surface | 3 |
| | | Small, scattered fissures or cracks | 2 |
| | | Several, small or few but large fissures | 1 |
| | | Total degeneration of grafted area | 0 |
| 4 | Color of repair tissue | Hyaline, pearly like | 4 |
| | | Predominantly hyaline, pearly like (>50%) | 3 |
| | | Predominantly white (>50%) | 2 |
| | | White | 1 |
| | | No repair tissue | 0 |
| Overall Repair Assessment | | Grade I | 16 |
| | | Grade II | 12 - 15 |
| | | Grade III | 8 - 11 |
| | | Grade IV | 4 - 7 |
| | | Grade V | 0 - 3 |

Table S2. Scoring criteria for Goebel score

| | Goebel | Points |
|---|---|---|

| | | | |
|---|---|---|---|
| 1 | **Color of the repair tissue** | Hyaline, pearly like | 4 |
| | | Predominantly hyaline, pearly like (>50%) | 3 |
| | | Predominantly white (>50%) | 2 |
| | | White | 1 |
| | | No repair tissue | 0 |
| 2 | **Presence of blood vessels in the repair tissue** | No blood vessels | 4 |
| | | Less than 25% of the repair tissue | 3 |
| | | 25-50% of the repair tissue | 2 |
| | | 50-75% of the repair tissue | 1 |
| | | More than 75% of the repair tissue | 0 |
| 3 | **Surface of the repair tissue** | Smooth, homogeneous | 4 |
| | | Smooth, heterogeneous | 3 |
| | | Fibrillated surface | 2 |
| | | Incomplete new repair tissue | 1 |
| | | No repair tissue | 0 |
| 4 | **Filling of the defect** | In level with adjacent cartilage | 4 |
| | | >50% repair of defect depth or hypertrophy | 3 |
| | | <50% repair of defect depth | 2 |
| | | 0% repair of defect depth | 1 |
| | | Subchondral bone damage | 0 |
| 5 | **Degeneration of adjacent articular cartilage** | Normal | 4 |
| | | Cracks and/or fibrillations in integration zone | 3 |
| | | Diffuse osteoarthritic changes | 2 |
| | | Extension of the defect into the adjacent cartilage | 1 |
| | | Subchondral bone damage | 0 |
| **Total points** | | | 20 |

Table S3. Histopathological scoring criteria

| | Parameter | Qualifications (%) | Points |
|---|---|---|---|
| 1 | **Defect Fill** | 100 | 4 |
| | | 75 - 99 | 3 |
| | | 50 - 74 | 2 |
| | | 25 - 49 | 1 |
| | | 0 - 24 | 0 |
| 2 | **Cell Distribution** | All | 4 |
| | | 67 - 99 | 3 |
| | | 34 - 66 | 2 |

| | | | |
|---|---|---|---|
| | | 1 - 33 | 1 |
| | | None | 0 |
| **3** | **Perilesional cloning** | None | 3 |
| | | Seldom | 2 |
| | | Occasional | 1 |
| | | Frequent | 0 |
| **4** | **Lesion-perilesion tissue integration** | Complete | 2 |
| | | Gap on one side | 1 |
| | | Gap on both sides | 0 |
| **5** | **Subchondral bone attachment** | 100 | 4 |
| | | 75 - 99 | 3 |
| | | 50 - 74 | 2 |
| | | 25 - 49 | 1 |
| | | 0 - 24 | 0 |
| **6** | **Surface fibrillation** | None | 3 |
| | | Slight fibrillation | 2 |
| | | Moderate fibrillation | 1 |
| | | Severe fibrillation | 0 |
| **7** | **tidemark reformation** | Complete | 3 |
| | | 51 - 99 | 2 |
| | | 1 - 50 | 1 |
| | | None | 0 |
| **8** | **Saf-O staining** | 91 - 100 | 4 |
| | | 76 - 90 | 3 |
| | | 51 - 75 | 2 |
| | | 26 - 50 | 1 |
| | | 0 - 25 | 0 |

Table S4. Subscores for modified ICRS gross scoring system. Means ± SD are reported.

| Modified ICRS Subscore Parameter | Control (n=8) | Saline (n=6) | rhLubricin (n=6) |
|---|---|---|---|
| **Degree of Defect Repair** | 4±0 | 2.8±0.8 | 3.1±0.4 |
| **Integration to Border Zone** | 4±0 | 2.2±0.8 | 2.6±0.4 |
| **Macroscopic Appearance** | 4±0 | 1.8±0.4 | 2.3±0.6 |
| **Color of Repair Tissue** | 4±0 | 2.3±0.3 | 2.4±0.7 |

Table S5. Subscores for Goebel gross scoring system. Mean ± SD are reported.

| Goebel Subscore Parameter | Control (n=8) | Saline (n=6) | rhLubricin (n=6) |
|---|---|---|---|
| **Color of Repair Tissue** | 4±0 | 2.3±0.3 | 2.4±0.7 |
| **Presence of Blood Vessels** | 4±0 | 2.1±1.4 | 2.8±0.6 |
| **Surface of Repair Tissue** | 4±0 | 2.1±0.2 | 2.3±0.6 |
| **Filling of Defect** | 4±0 | 2.8±0.6 | 3.3±0.3 |
| **Degeneration of Adjacent Cartilage** | 4±0 | 2.4±0.6 | 2.8±0.4 |

Figure S1. Safranin-O sections for all samples used in histological assessments (n=3/group). Scale bars = 1 mm.

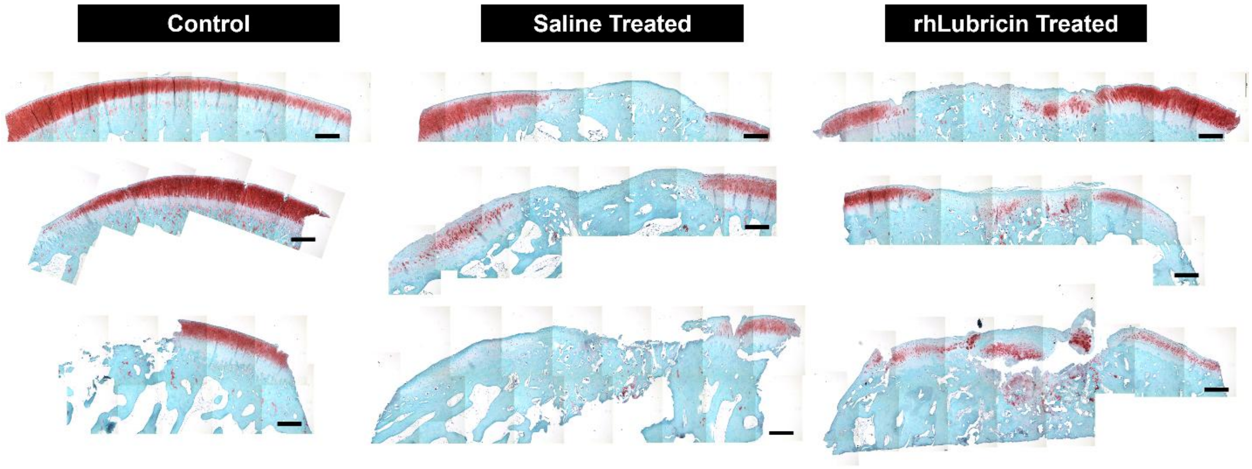
